# NudF-boosted strategy to improve the yield of DXS pathway

**DOI:** 10.1101/2022.03.10.483762

**Authors:** Devi Prasanna, Shaza Wagiealla Shantier, Ashish Runthala

## Abstract

**Background:** Terpenoids form a large pool of highly diverse organic compounds possessing several economically important properties, including nutritional, aromatic, and pharmacological properties. The DXP pathway’s end enzyme, nuclear distribution protein (NudF), interacting with isopentenyl pyrophosphate (IPP) and dimethylallyl pyrophosphate (DMAPP), is critical for the synthesis of isoprenol/prenol/downstream compounds. The enzyme is yet to be thoroughly investigated to increase the overall yield of terpenoids in the *Bacillus subtilis*, which is widely used in industry and is generally regarded as safe (GRAS) bacterium. The study aims to analyze the evolutionary conservation across the active site, and map the key residues for mutagenesis studies. The study would allow us customize the metabolic load towards the synthesis of prenol or isoprenol or any of the downstream molecules.

**Results:** The 37-sequence dataset, extracted from 103 Bacillus subtilis entries, show a high phylogenetic divergence, and only six one-motif sequences ASB92783.1, ASB69297.1, ASB56714.1, AOR97677.1, AOL97023.1, and OAZ71765.1 show monophyly relationship, unlike a complete polyphyly relationship between the other 31 three-motif sequences. Further, only 47 of 179 residues of the representative sequence CUB50584.1 are observed to be significantly conserved. Docking analysis shows a preferential bias of ADP-ribose pyrophosphatase towards IPP, and a nearly 3-fold energetic difference is observed between IPP and DMAPP. Computational saturation mutagenesis of the seven hotspot residues identifies two key positions LYS78 and PHE116, encoded within loop1 and loop7, majorly interact with the ligands DMAPP and IPP, and their mutants K78I/K78L and PHE116D/PHE116E are found to stabilize the overall conformation. The loops are hereby shown to play a regulatory role in guiding the promiscuity of NudF towards a specific ligand.

**Conclusion:** The study map the phylogenetic relationship between the 37 representative *B*.*subtiis* NudF sequences, and through sequence conservation, structural contact map, topological flexibility, and saturation mutagenesis of the active site residues, the essential residues regulating the interaction of NudF with IPP/DMAPP are deciphered. The study robustly screens its mutational landscape and localizes the two crucial residues LYS78 and PHE116 for directing the mutagenesis studies. The preliminary docking and simulation results also suggest a preferential bias of ADP-ribose pyrophosphatase towards IPP over DMAPP. The findings would pave the way for the development of novel enzyme variants with highly improved catalytic ability for the large-scale bioproduction of specific terpenoids with significant neutraceutical or commercial value.

## 1. Introduction

Isoprenoids are the most functionally and structurally varied class of secondary metabolites, with over 55,000 identified molecules (Vickers et al. 2017; Christianson, 2008) and have been used to make aromatic, flavoring, and medicinal molecules (Phan-Thi et al. 2019; Phulara SC*et al*. 2020; Matulja et al. 2020). As the natural extraction of isoprenoid based bioactives has led to an overexploitation of plants, the global research interest has now shifted to utilize the generally regarded as safe (GRAS) status microbes like *Bacillus subtilis* as the biocatalytic machinery (Zhou et al. 2013). A key protein controlling the yield of an end-product molecule is ADP-ribose pyrophosphatase or NudF (EC 3.6.1.13) that orderly hydrolyzes DMAPP and IPP to prenol and isoprenol (Figure1), collectively termed as isopentenol (C_5_ alcohol), a building block of all the higher-order downstream terpenoid molecules (Formighieri and Melis, 2014). It is a member of the Nudix superfamily (Pfam PF00293; InterPro IPR000086) (McLennan, 2006), and plays a key role in the synthesis of AMP and ribose-5-phosphate from ADP-ribose, actively dephosphorylating the phosphate moieties of several diverse substrates with a broad versatility. Its close homolog is the NudB protein (EC:3.6.1.67) from *E. coli* (EcNudB), having a catalytic activity towards GPP and FPP (Mildvan et al. 2005). It has been suggested that an additional phosphatase AphA is needed to hydrolyze IPP to isoprenol because NudB can only catalyze the hydrolysis of IPP to isopentenol (Tian et al. 2019). While EcNudB exhibits a strong affinity for DMAPP, the NudF protein of *Bacillus subtilis* (BsNudF) exhibits no such preference, and equally interacts with both DMAPP and IPP to orderly produce an equivalent amount of prenol and isoprenol (Withers et al. 2007). A varying affinity for various substrates may substantially increase pressure on the stoichiometric flux, and until the connected regulatory network of proteins does not drain out the added molecules at an equivalent rate, it becomes toxic to the cell and inhibits cell growth. However, both these enzymes show a low affinity towards IPP/DMAPP. Hence, to escape this major bottleneck, a fusion protein of Isopentenyl-diphosphate delta isomerase (IDI), also known as Isopentenyl pyrophosphate isomerase (IPP isomerase), and EcNudB has been utilized and with an increased expression, the metabolic flux increases the bioproduction rate of only prenol and reduces the isoprenol production. As an attempt to increase the production rate of downstream molecules in *E*.*coli*, an endogenous overexpression of IspG and 1-deoxy-D-xylulose-5-phosphate synthase(DXS) and exogenous expression of YhfR and BsNudF has been used (Liu et al. 2014).

**Figure 1:**
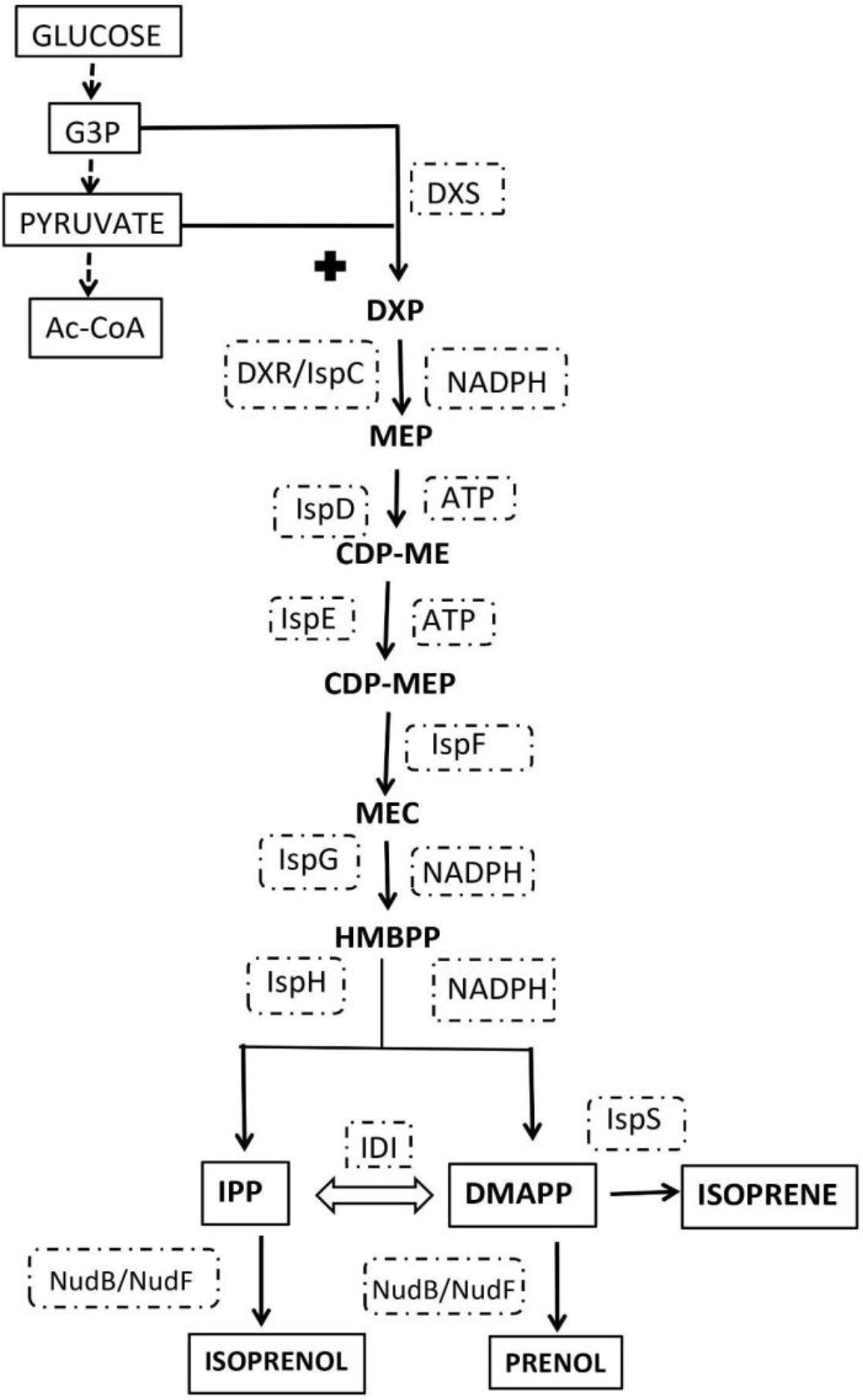
The MEP pathway, representing the biocatalysts at all intermediary stages. From the glyceradehyde-3-phosphate (G3P) and pyruvate, it synthesizes the 5-carbon building blocks DMAPP and IPP for producing terpenoids. It includes an ordered set of 7 enzymes viz. DXS, DXP reductoisomerase (DXR),2-C-methyl-D-erythritol 4-phosphate cytidylyltransferase (IspD), 4-diphosphocytidyl-2-C-methyl-D-erythritol kinase (IspE), 2-C-methyl-D-erythritol 2,4-cyclodiphosphate synthase (IspF), HMB-PP synthase (IspG) and HMB-PP reductase (IspH), followed by the interplay of IDI and NudF.

Although sufficient IPP and DMAPP concentrations are required for terpenoid synthesis (Formighieri and Melis, 2014), an excess of these molecules can inhibit cell growth (Withers et al. 2007) and thus reduce terpenoid production (Sivy et al. 2011), and therefore NudF enzyme continuously use these potentially harmful debris molecules. To increase the yield of DXS pathway, researchers have specifically targeted the first rate-limiting enzyme DXS (Lange et al. 1998; Estevez et al. 2001) because its K_cat_/K_m_ score is much smaller than all the other enzymes (Kuzuyama et al. 2000). Improving the DXS activity has been considered as the most effective measure to improve the overall yield for several species, including *Streptomyces* (Kuzuyama et al. 1998), *Lycopersicon esculentum* (Rohmer 1999) and *Synechococcus leopoliensis* (Schwender et al. 1996; Bach and Lichtenthaler 2010). Heterologous expression of *Bacillus subtilis* DXS in *E*.*coli* (Leonard et al. 2010) has also been shown to increase the overall yield. Yet another strategy, using the functionally mutated recombinant poplar DXS (Banerjee et al. 2016), has shown a negligible feedback inhibition for IPP/DMAPP and has been fruitful. To mitigate the toxicity of IPP, the farnesyl diphosphate synthase and IDI synthase have been overexpressed in *E*.*coli*, and it has led to an 800-fold production of sesquiterpene β-farnesene (You et al. 2017). Likewise, the balanced metabolic concentration of IPP and DMAPP (Hahn etal. 1999; Yoon etal. 2007), heterologous expression of*Haematococcus pluvialis* IDI (Sun et al. 1998), or overexpression of *Lycium chinense* IDI in *E*.*coli*(Li et al. 2016) have been shown to increase the overall yield. Although the majority of these methods have overexpressed the rate-limiting enzymes, it may lead to a complete metabolic imbalance and significantly minimize the yield of downstream terpenoids. In this regard, directed co-evolution of DXS, DXR and IDI have been shown to increase isoprene production (Li et al. 2016). However, these strategies have also not efficiently focused the major enzyme BsNudF for increasing the overall yield, and directed evolution of NudF‟s promiscuous active site would therefore be significantly useful to increase the biosynthetic rate of a downstream terpenoid. For understanding the preferential binding of IPP/DMAPP, catalytic and behavioral switching of BsNudF, this article deciphers the key functional details across the active site for decoding the mutations that could improve its activity. As it comprehensively maps the crucial residues at the functionally important positions, the study will be fruitful in designing a custom set of key interacting residues against a required ligand, to attain a theoretically impossible overall yield. Although DXS shows the Km score of∼96 for pyruvate and ∼240 for pyruvate, and Kcat score of∼270 (Kuzuyama et al. 2000), NudF is pivotal for successfully improving the biocatalytic yield of any downstream terpenoid, the limited molecular engineering strategy, especially focused on it, is of prime interest for biological and industrial research to optimally escape the bottleneck and channel the biosynthetic yield towards the production of the desired end-product.

## 2. Theoretical Calculations

### 2.1 Construction of sequence dataset and functional affirmation

Screening the NudF protein sequences in the NCBI Protein database, a set of 105,910 protein sequences is found, of which only 103 *Bacillus subtilis* entries are found and are used to build the primary dataset. Purging the completely redundant entries with at least 80% alignment coverage through MMSeqs2, a reduced dataset of 40 entries is derived, and their length pattern is noted. Three entries WP_139026580.1, WP_139026569.1, CUB36584.1 orderly encoding only 118, 86, and 55 residues, are found to be the partial sequences and are purged. A final dataset of 37 entries, including 1, 6, and 30 sequences orderly encoding 179, 205, and 185 residues, is thus created. Before pursuing the analysis further, it is mandatory to functionally affirm the defined dataset. The sequences are fed to MEME (Bailey et al. 2009) for screening the conservation of the signature motif, and affirming the functions of the sequences to use the functionally correct entries. As no functionally similar *B*.*subtilis* protein has been structurally resolved to date, the homologous *Escherichia coli* structure (PDB ID: 5U7E) is used as the representative entry to reliably localize the conserved motifs. Further, to deploy the *Bacillus subtilis* representative sequence for the downstream analysis, the smallest sequence (CUB50584.1) is used from the constructed dataset. It is because evolution tends to decrease a protein sequence length to pack the function more optimally (Lipman et al. 2002).

### 2.2 Modelling the representative sequence

As the *B*.*subtilis* representative sequence CUB50584.1 is still not experimentally determined, its tertiary structure is modelled through template-based modelling methodology using MODELLER, as per the recently published strategy (Runthala 2021; Runthala and Chowdhury, 2019). For reliably predicting the model, the wild-type nucleoside diphosphate sugar hydrolase from *Bdellovibrio bacteriovorus* (PDB ID: 5C7Q) is used as the template, sharing a sequence identity of 36%. As the first predicted protein model usually has several non-physical local clashes, the constructed model is energetically relaxed through the conservative refinement strategy of Galaxyrefine2 to maximally retain the topology extracted from the template. The refined model is subsequently evaluated through Swissmodel (Benkert et al. 2011) to assess its credibility.

### 2.3 Phylogenetic and conservation analysis

Statistically significant evolutionary relationships are typically considered for drawing the meaningful phylogenetic connections within the selected set of sequences/species. As correct sequence alignment is certainly required for extracting the accurate phylogenetic relationship, the 37-sequence *B*.*subtilis* dataset is aligned through the hidden markov model based clustal-omega module of HHPred (Zimmermann et al. 2017). The constructed profile is fed to the IQTree server (Trifinopoulos et.al. 2016) for deriving the evolutionary tree on the basis of the default parameters, using the default ultrafast methodology over 10000 bootstrap alignments at the minimum correlation coefficient or the convergence threshold of 0.99. The resultant consensus tree is visualized and analyzed through ITOL (Letunic and Bork, 2019). Further, the sequence alignment is fed to Consurf (Ashkenazy et.al. 2016) for plotting the average sequence conservation scores over the predicted CUB50584.1 model, using the default conditions, and analyzing the sequence variations across the functionally crucial sites. Lastly, the constructed alignment is also visualized using Espript (Gouet et.al. 2003) to mark the structural conservation.

### 2.4. Active site prediction and docking analysis with IPP and DMAPP

To reliably channelize the interaction of substrates only at the biologically credible site(s) and to exclude any fake docking solution, the CastP version3 server is used to predict the active site(s) (Tian et al. 2018). For screening the most promising energetically feasible interaction site(s) of NudF against the substrates IPP and DMAPP, the Dockthor server (https://www.dockthor.lncc.br/v2/) is used (Guedes et al. 2021), using the default parameters. For the six and five rotatable bonds of these ligands, a complete degree of rotational freedom is orderly allowed to attain biologically correct predictions. To enhance the accuracy, the server specifically improves the torsional energy score for the ligand on basis of its conformational flexibility, and further including the energy components of lipophilic contacts, polar and non-polar solvation, it redefines the overall energy function of the MMFF94S force field through machine learning. The resultant solutions are subsequently analyzed using UCSF Chimera to extract the key interacting residues.

### 2.5 Crucial residues for functional mutagenesis

The accuracy of a rationally evolved enzyme molecule is dependent on the identification of the hotspot residues, whose mutations could enhance its catalytic activity (Hu et al. 2020). To analyze the hotspot regions, proximal to the active site and tunnel, and appropriately map the mutational landscape of the predicted protein molecule, the hotspot server 3.1 (Sumbalova et al. 2018) is used. Although this server is capable of modelling an input protein sequence, the constructed CUB50584.1 model is considered to drive the analysis for the topologically reasonable model. As this server also robustly estimates the thermodynamic stability of a mutation through FoldX and Rosetta, it is shown to reliably exclude the destabilizing mutations (Pucci et al. 2015), shown to majorly decrease the prediction quality (Pucci et al. 2018). However, to still extract its fullest potential and reliably select the best mutations from the preselected resultant ones, the resulting data is analyzed along with the other residues marked in the preceding steps.

To understand the most promising mutations across the active site, the correlated mutations are mapped for the 179residue-sequence CUB50584.1 sequence through the GREMLIN server (Ovchinnikov et al. 2014). As these coevolution-based contacts are crucial to reliably discriminate the true hotspot residues from the spurious ones, the contact map network is analyzed to mark the true hotspots. Purging the unreliable positions, the true hotspots are analyzed through the Dynamut2 server (Rodrigues et al. 2021), trained, and tested on the Protherm database (Kumar et al. 2006). Its high accuracy is evident from the fact that it outperforms the other measures with a Pearson‟s correlation of up to 0.72 and 0.64 for the single and multiple point mutations with an RMSE error (Kcal/mol) of 1.02 and 1.8 respectively, making it a trustworthy strategy for prioritizing the stabilizing/destabilizing mutations (Rodrigues et al. 2021).

### 2.6 Computational Mutagenesis and flexibility analysis

On the basis of docking and mutagenesis results, the most favorable binding and substrate preferences are excavated. To robustly quantify the strength of interaction for each of the ligands, the Dynamut2 results are analyzed for the five prioritized residues (ARG18, ALA117, ASP139, GLU140, ASP141), and their five top-ranking ΔΔG scores are analyzed. To examine this data, NetsurfP2.0 (Klausen et al. 2019), with an 80 percent correlation with experimentally confirmed data, is preferred to estimate the relative solvent accessible surface area (RSA) for the five key positions for the native protein. For further assessing the flexibility across the NudF structure, the flexibility of its Cɑ-backbone is predicted using the CABS-flex 2.0 which computes an average topological fluctuation-diversity of 10 medoids of the 10 cluster sets, derived from a 10ns-simulated set of 1000 decoys (Kuriata et al. 2018). To further get more insights of the binding affinity of the two ligands, a small 100-ns molecular dynamics (MD) simulation is finally done to analyze the physical movements of ligand and protein atoms. The IPP and DMAPP topologies are constructed through PRODRG2 (Schüttelkopf and van Aalten, 2004), and CUB50584.1 model and its two complexes are simulated through the WebGro server (https://simlab.uams.edu/).

Utilizing the GROMACS96 43a1 forcefield, SPC water model and triclinic simulation box type, the structures are simulated by neutralizing its charge with 0.15M sodium chloride. Energetically refining the structures through the steepest descent function, integrated at every 5000 steps, the structures are subjected to NVT/NPT equilibrations at the default conditions of 300K temperature and 1bar pressure. A short 100-ns MD simulation is implemented by using 1000 frames/simulation to integrate the trajectory through leap-frog algorithm for lastly assessing the resultant trajectory through all of its encoded parameters, viz. time-dependent root-mean-square deviation (RMSD) for the overall structure and RMS fluctuations (RMSF) across its residues, radius of gyration (Rg), number of hydrogen bonds, overall and per-residue solvent accessible surface area (SASA) and ligand-RMSD.

## 3. RESULTS AND DISCUSSIONS

### 3.1 Construction of sequence dataset and its functional affirmation

To affirm the functional attribute of all the derived 37 sequences, the relative location of their three top-ranked conserved motifs is screened against the functionally deciphered structure (5U7E) of *E*.*coli* through MEME (Figure 2A). The members of the nudix superfamily encode the signature sequence GX_5_EX_7_REUXEEXG/TU, where U is either L/V or I (Bessman et al. 1996) as represented in green in this figure. This conserved sequence is responsible for metal-binding and forms the catalytic site in more than 4000 enzymes in numerous species including eukaryotes, prokaryotes, and viruses (Gabelli et al. 2007). The NudF length range is shown to be 179-205, with 182-residue sequence being the most common. Here, the six 205-residue sequence subset shows a stark feature, i.e. ASB92783.1, ASB69297.1, ASB56714.1, AOR97677.1, AOL97023.1, and OAZ71765.1encode only one characteristic motif, as observed for 5U7E, unlike the other *B*.*subtilis* sequences having all the three conserved motifs.

**Figure 2:**
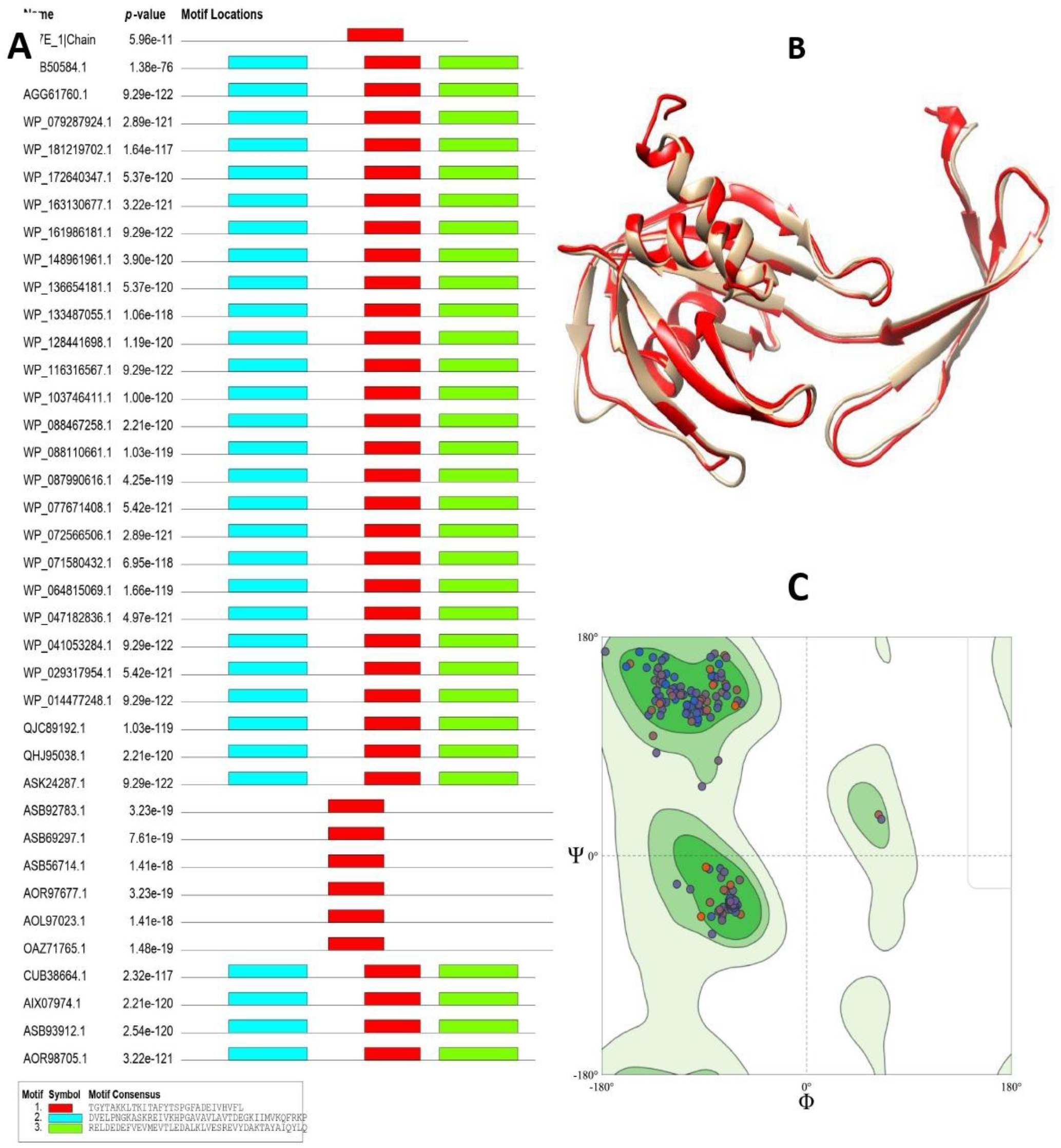
(A) Top-3 motifs, screened against the functionally deciphered structure 5U7E of *E*.*coli*, (B) Structural overlap of the constructed model over 5U7E (C) Ramachandran map of the predicted model.

### 3.2 Modelling the representative sequence

The representative *B*.*subtilis* sequence CUB50584.1 is modelled using 5C7Q through the template-based modelling methodology using MODELLER9.25. The predicted structure is refined using GalaxyRefine2 and is subsequently evaluated through Swissmodel (Benkert et al. 2011). The model shows an RMSD score of 0.887 against the deployed template (Figure2B). The model shows a Molprobity score of 1.28, indicating a topological accuracy of a reasonably high accuracy X-ray structure. Further, 98.87% of residues are found localized within the Ramachandran favored regions (Figure2C).

### 3.3 Phylogenetic and conservation analysis

Feeding the sequence alignment of HHPred-clustal omega to IQTree, and building the evolutionary tree using 10000 bootstrap alignments at the convergence threshold of 0.99, the resultant consensus solution is visualized and analyzed through ITOL (Figure 3). Although the single motif-six sequence subset (AOR97677.1, OAZ71765.1, ASB69297.1, ASB92783.1, ASB56714.1, and AOL97023.1) appears to be clustered with 5U7EA and the representative sequence CUB50584.1, encoding all three motifs, their average sequence identity is 30.3640.339, compared to 14.73 and 21.3630.3304 against 5U7EA and CUB50584.1, respectively. Moreover, all the other sequences are not found to share substantial sequence similarity, as also observed with their scrutiny through Clustal Omega (Zimmermann et al. 2017). All the other sequences are found uncluttered with any other entry, and as recently shown in the phylogenetic study of 80000 Nudix homologs, a general monophyly is prominent, besides a few occasional incidences of homoplasy (Srouji et al. 2017). The constructed profile is further fed to Consurf for mapping the average sequence conservation scores over the predicted structure, using the default conditions (Ashkenazy et.al. 2016), and the sequence variation across the functionally crucial sites is analyzed (Figure 4). A set of 14 residues, viz.ILE20, ALA45, GLN62, GLY77, GLU80, GLY82, ALA89, GLU92, GLU95, GLU96, THR112, ALA126, THR130, and GLU148 are found to be completely conserved, with a conservation score of 9. Moreover, 33 residues LEU4, THR8, PHE15, PRO30, ASN31, LYS36, ILE39, HIS42, PRO43, ALA50, LYS56, VAL60, LYS65, ILE71, PRO85, THR88, ARG91, LEU93, GLU94, THR97, LEU106, ILE107, GLU119, TYR124, ASP141, VAL144, ALA154, LEU157, ASP166, LYS167, THR168, PHE170 and GLN173 show a conservation score of 7 and 8, indicating significantly higher conservation. Further, 11 residues, viz. PRO13, ARG23, VAL24, ALA33, MET34, LYS69, LYS131, SER150, GLU153, ASP160, and HIS164 are found to be completely variable across the defined dataset. It is astonishing that the representative *B*.*subtilis* sequence encodes mere 11 (6.145%) variant residue loci in contrast to the statistically higher sequence conservation at 47 (26.259%) positions (Figure4), and still the enzyme is able to show a highly promiscuous nature, and it indicates that a few key residues should play a major role within the active site of this enzyme.

**Figure 3:**
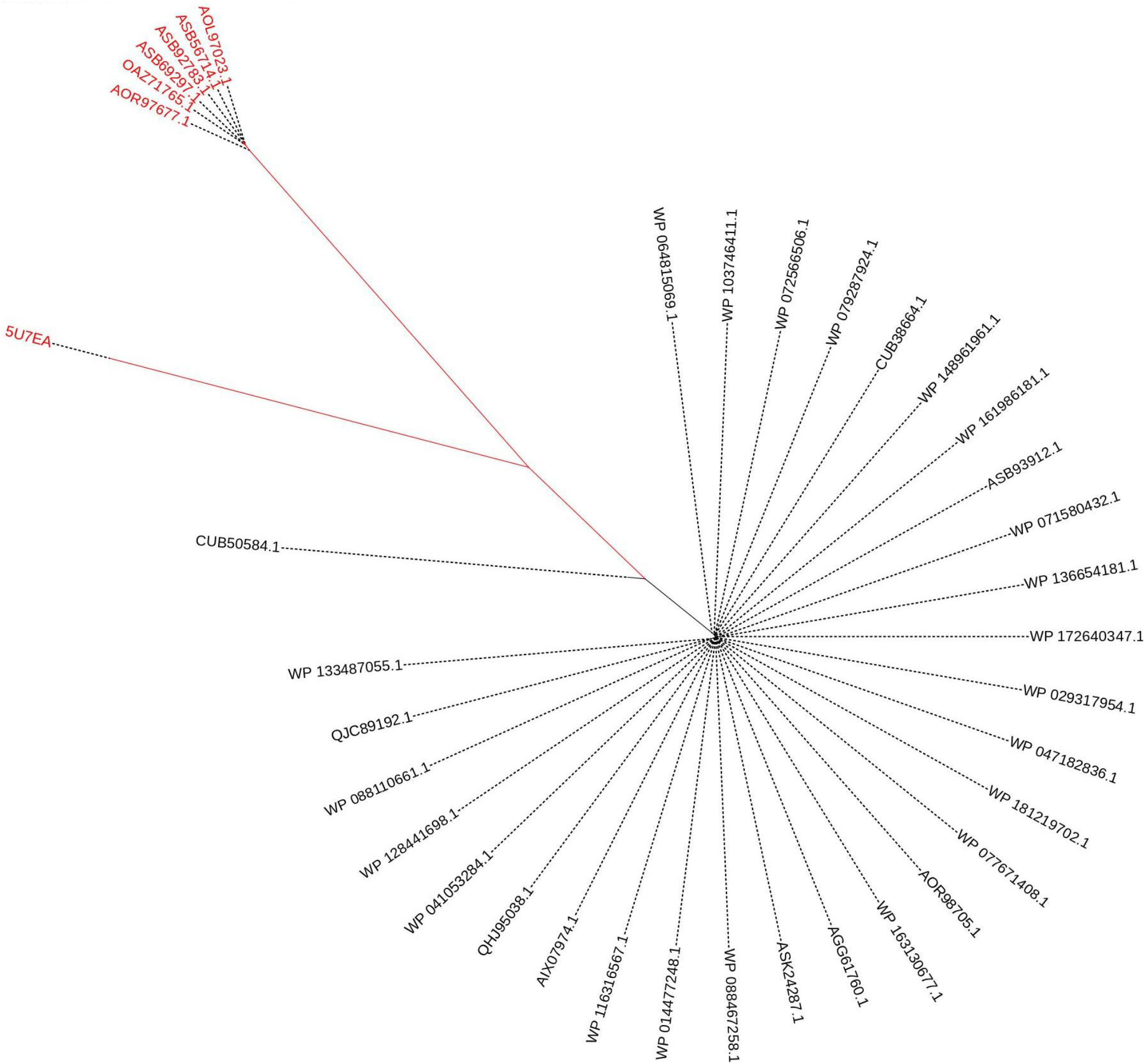
Unrooted circular tree of the 37-sequence NudF dataset.

**Figure 4:**
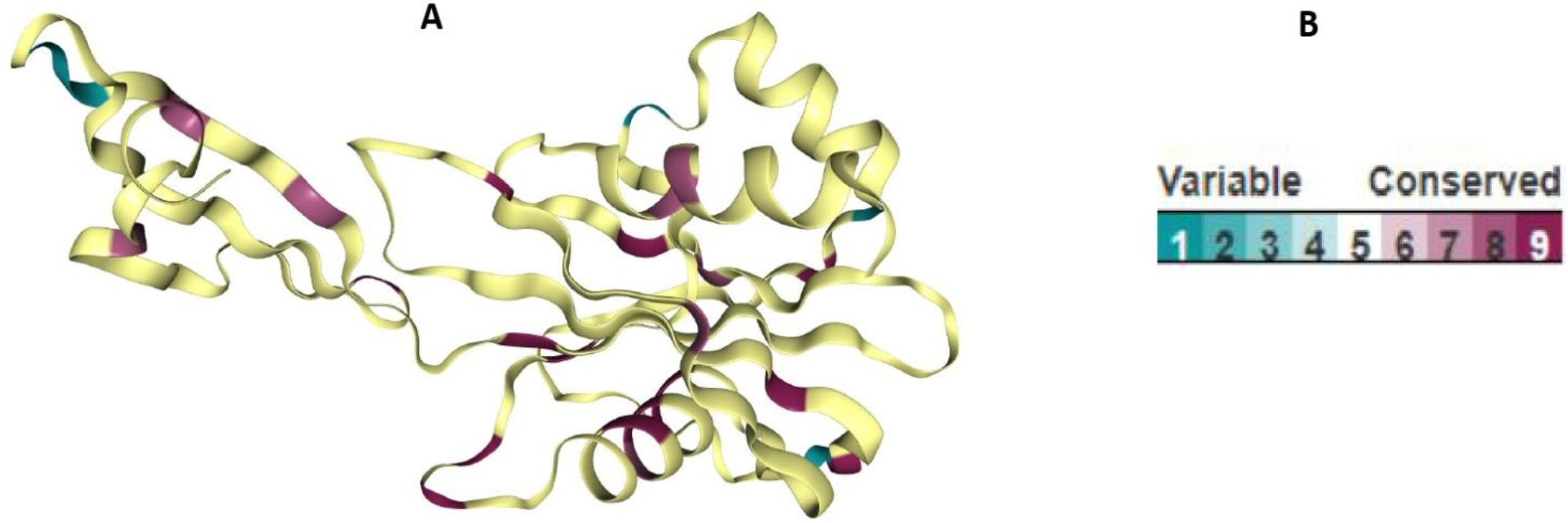
(A) Consurf-derived sequence conservation for the 37-sequence NudF dataset, projected onto the representative sequence model CUB50584.1 (B) Color range varying between 1 and 9, maroon referring to the completely conserved positions.

### 3.4 Active site prediction and docking analysis with IPP and DMAPP

Excluding the shallow openings and using a probe radius of 1.4Å, CastP (Tian et al. 2018) delineates the empty concavities on a protein surface to map the volume spectrum of cavities and pockets. It robustly deciphers the surface properties and localizes the functionally important zone, and shows only 2 biologically meaningful active pockets in the predicted NudF model, with molecular surface area (Å^2^) and molecular volume (Å^3^) of 501 and 946.6 (Pocket1), and 208.9 and 661.3 (Pocket2) respectively, and this is in accordance with the known structural details of the NudF‟s *E*.*coli* homolog, wherein two active sites have been observed (Gabelli et al. 2002). Further, these pockets are orderly found to span a set 12 (MET1, LEU4-A-GLU6, ARG37, ILE39, TYR111, PRO114, GLY115, ALA117, ASP118, ILE120) and 30 (ARG18-VI-LYS21, VAL40-NH-PRO43, ALA45, VAL60, GLN62-YRK-ALA66, GLU73-IPAG-LYS78, GLU96, THR112, GLU119, LEU121, ASP141, GLU142, ASP165, ALA166 and LYS167) residues. Here, it is worth noting that Dockthor uses only one most-voluminous docking site for both ligands (Pocket1, Figure5A), and it indicates a preferential binding of ligands over the second superficial site (Pocket2, Figure5B).

As a modest level of NudF is enough to overcome the IPP/DMAPP toxicity (Withers et al. 2007), this enzyme must interact with both of these substrates, and this responsible molecular NudF surface (Figure5C) closely interacts with the two ligands (Figure5D). The ligands IPP and DMAPP orderly show an interaction energy (Kcal/mol) and affinity score (Kcal/mol) of - 115.388 and −6.431, and −41.402 and −5.271. Screening the active site residues within 5Å of these two ligands, it is observed that 14 residues viz. VAL19, ILE20, HIS42, ALA45, GLN62, ARG64, GLU73, ALA76, LYS78, THR112, GLU119, LEU121, ASP165, LYS167, and 8 residues viz. VAL22, GLU38, VAL40, HIS42, SER113, PHE116, ALA117, and GLU119 orderly interact with the two ligands IPP(yellow) and DMAPP(blue). To analyze it further, the two-dimensional and three-dimensional interaction maps are drawn for DMAPP (Figure 5E and F) and IPP (Figure 5G and H). It indicates that two key hotspot residues LYS78 and PHE116, orderly responsible for interacting with these ligands through one and two residues in the active site, could be the key to specifically alter the active site to stabilize its affinity for the required ligand.

**Figure 5:**
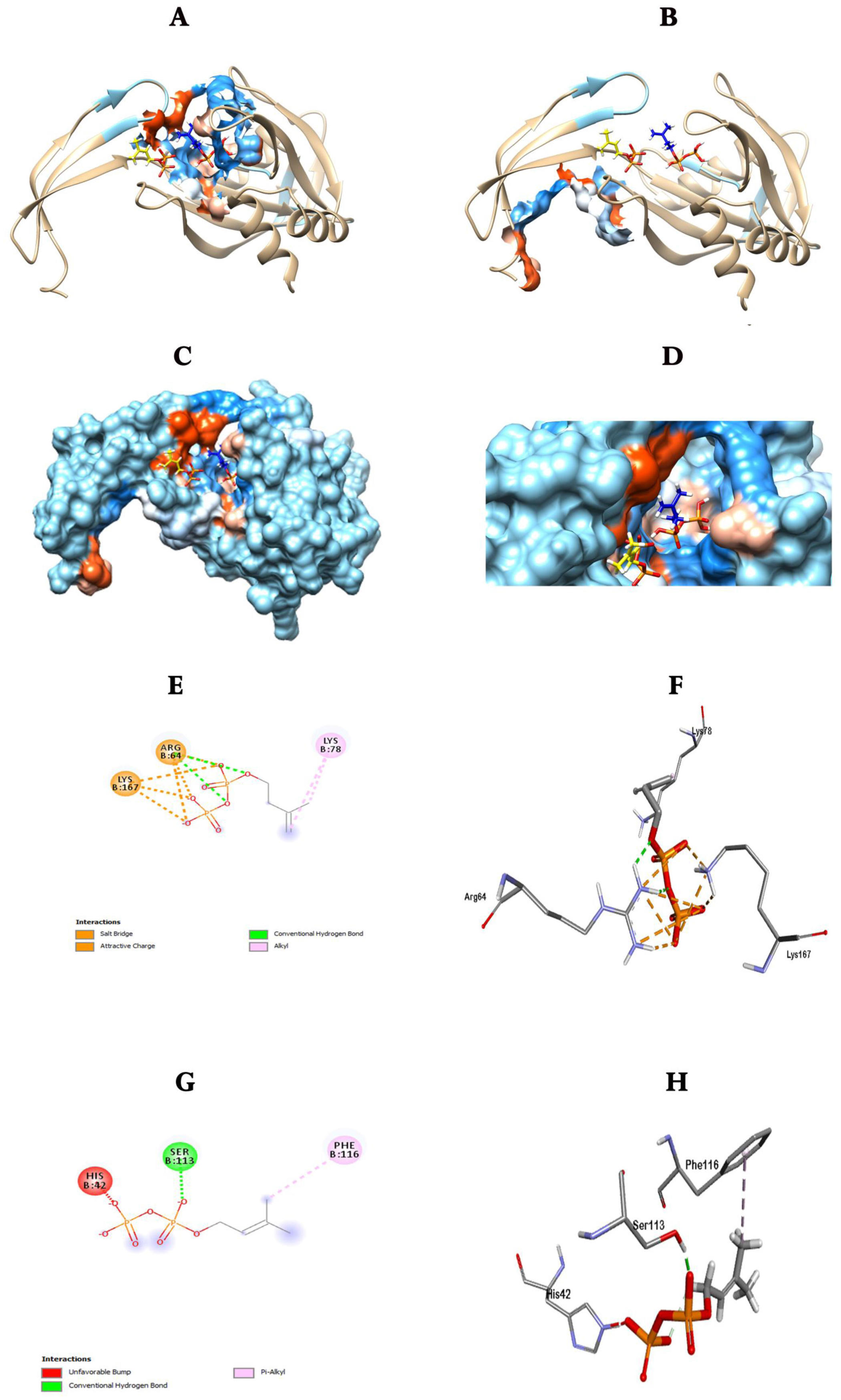
Dockthor results of both IPP and DMAPP. (A) Pocket1 (B) Pocket2 represented in correlation with the two ligands IPP(yellow) and DMAPP(blue), along with the close analytical view of Pocket1, in terms of (C) Molecular surface (D) Active site topology at the vent of the tunnel. (E) Two-dimensional (F) Three-dimensional interaction maps of protein with DMAPP, (G) Two-dimensional (F) Three-dimensional interaction maps of protein with IPP show the key residues interacting within the active site.

### 3.5 Crucial residues for functional mutagenesis

Through GREMLIN (Ovchinnikov et al. 2014), the contact map network of the modelled protein is constructed. For all the predicted contacts, it yields the two-dimensional distance matrix along with their probability scores, and is shown to be accurately predicting both direct and indirect residue couplings. The coevolution-based network for CUB50584.1 is analyzed (Figure 6A), and the statistically top-ranked contacts are extracted. A set of 22 functional hotspot residues, viz. ARG18, ILE20, LYS21, ASN41, PRO43, LEU67, ILE71, LYS78, LEU79, PRO81, GLY82, PHE110, TYR111, THR112, PHE116, ALA117, LEU121, ASP139, GLU140, ASP141, ALA166, and PHE170, with a theoretically credible probability score higher than 0.5, are found in NudF. However, LEU79 is found to be completely buried in the structural core of NudF. Moreover, ILE20 and THR112 are found to be completely conserved and are thus excluded. The 19 key hotspot residues are found to define a structural network of 0-4 contacts. Plotting the number of contacts for these hotspots (Figure6B), seven residues, viz. ARG18, PRO43, PRO81, ALA117, ASP139, GLU140, and ASP141 are not found to have any contacts. As the delicate balance between the flexibility and stiffness, sustained by key contacts, majorly regulate its functional role(s), these seven residues (Figure7) are selected for further analysis. However, as loop PRO43 and PRO81 usually stabilize a structural bend through internal hydrogen-bonding, as reported earlier (Li et al. 1997, Trevino et al. 2007), these positions are also not considered for subsequent computational mutagenesis study.

**Figure 6:**
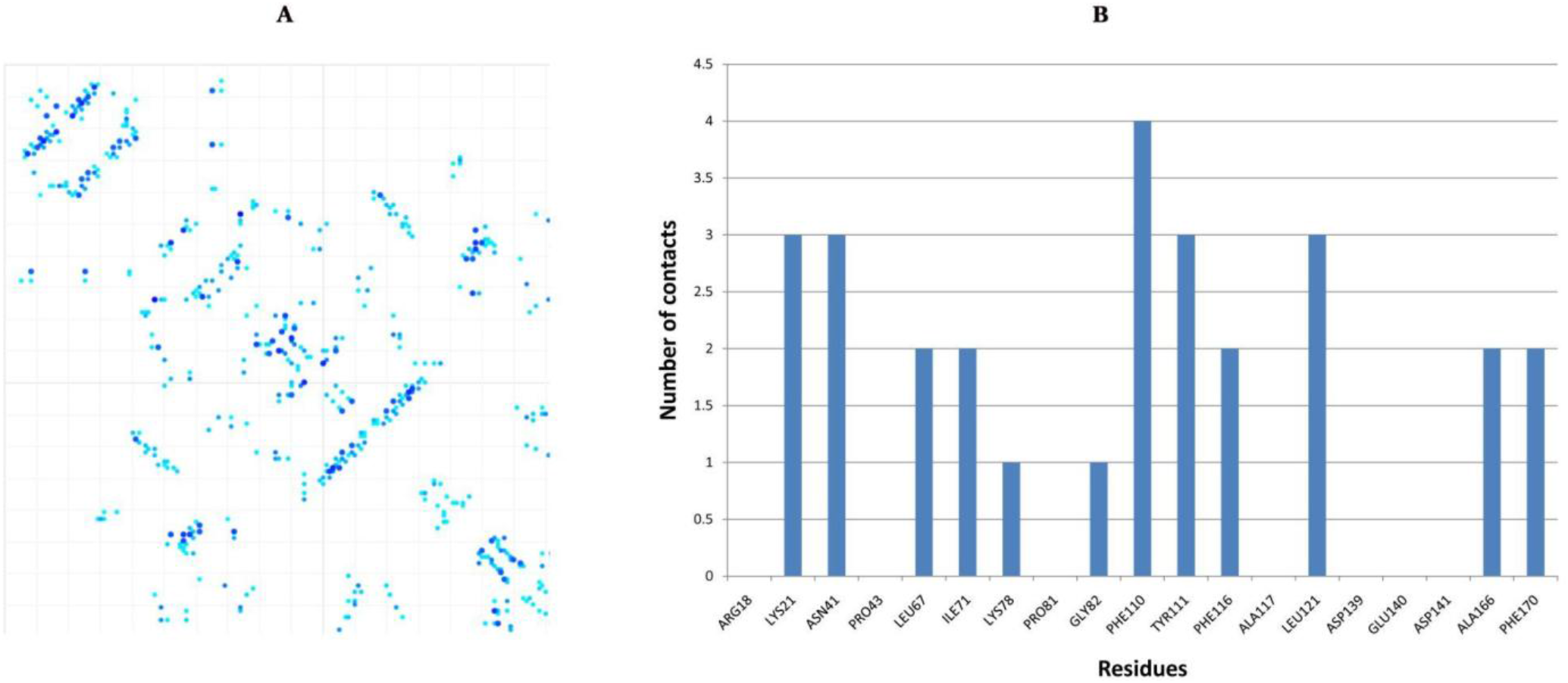
GREMLIN resultant (A) Contact map network (B) Number of contacts for the 19 key hotspot residues, placed proximal to the active site

### 3.6 Computational Mutagenesis and flexibility analysis

To reliably estimate the stability changes upon a point mutation on the constructed CUB50584.1 model, and as per the excavated crucial regions, the Dynamut2 server is used to estimate the stability changes upon a point mutation. The method derives the scores through the topological environment property and dynamic behavior of a residue, and is recently shown to outperform the prediction measures including FoldX and MAESTRO (Rodrigues et al. 2021). Restricting the search to the five prioritized residues (ARG18, ALA117, ASP139, GLU140, ASP141), and the two hotspot residues interacting the most stabilizing mutations are selected. The ΔΔG scores (KJ/mol) of these mutations ranges between −1.67 to 1.06. To correctly drive the analysis, the relative surface accessibility (RSA) for the five selected residues is computed using NetSurfP2.0 (Klausen et al. 2019), and the prioritized residues orderly show an RSA score of 0.393, 0.232, 0.536, 0.696, 0.622, 0.449 and 0.397, categorized as exposed or buried at the threshold of 0.25.

To excavate the active site and its key catalytic residues, the average structural fluctuations across NudF are predicted using CABS-flex 2 (Kuriata et al. 2018). and a set of 8 loops: loop1-loop8, spanning from ILE14-ARG23, ASP26-ARG37, ILE39-VAL46, ALA50-VAL58, GL62-ILE72, GLY77-PRO85, TYR111-LEU121 and THR130-VAL144 are found to be flexible. These loops, encompassing a 48.6% structure, majorly define the activity site topology, and are envisaged to substantially regulate the functional nature of NudF. It is interesting to observe that the selected five residues ARG18, ALA117, ASP139, GLU140, ASP141 are encoded by the loop1, loop7 and loop8, and it could be there that the loop1 acts as a capping loop and loop7 and loop8 make the active site sufficiently voluminous to interact with IPP and DMAPP. It is similar to *E*.*coli* NudF loop9, which is stabilized by its closed conformation (Gabelli et al. 2002).

To further analyze the dynamic atomic interactions of IPP/DMAPP within the active site, 100ns NudF and its complexes are simulated using WebGro, as shown recently(Vishvakarma et al. 2021, Tumskiy and Tumskaia 2021). MD simulation is a robustly accurate strategy to analyze the configurational changes that occur when a ligand is induced to fit (Leach, 2007). Using Webgro, the protein/complex system is computationally evolved using classical mechanics for a short 100ns timespan, and the configurational stability or binding affinity of a ligand is assessed across the simulation trajectory. To analyze the simulation results, Rg or the average distance between the center of mass and the rotational axis is usually used to estimate the conformational stability of a system against any physicochemical strain (Lobanov et al. 2008), and its lower score implies a higher stability. Here, the apoprotein shows the Rg scoring variations between ∼1.5-2.75nm, its complex with DMAPP and IPP orderly shows the respective scores of ∼1.8-2.05nm and ∼1.875-2.05nm, indicating that the apoprotein is relatively unstable than its complex structure, exactly like *E*.*coli* NudF (Gabelli et al. 2002). Moreover, IPP-complex is more stable than the DXP-complex because its trajectory mostly crawls at substantially lower scores across the trajectory. RMSD is another helpful measure for estimating the structural stability and the overall deviation from the backbone topology during at the complex formation at a specified temperature, as used earlier by Gromacs (Lindahl et al. 2010, Abraham et al. 2015). The backbone-based RMSD trajectory score should be the lowest because it directly gives the deviation of mean atomic coordinates and reflects the macromolecule’s conformational stability. Scrutiny of RMSD across the trajectory shows that the RMSD variations of apoprotein range between the acceptable range of ∼0.5-2.5nm in contrast to the respective ranges of 0.3-0.9 and 0.3-0.75 for its DMAPP and IPP complexes. Unlike the DMAPP complex trajectory, showing substantially higher undulations, the IPP complex gets stabilized after ∼60ns.

Furthermore, the RMSF, or residue fluctuations across the simulation, reveals a macromolecular system’s conformational stability, and its scoring undulations describe structural complexity, with a lower value signifying higher overall stability (Lindahl et al. 2010, Abraham et al. 2015). For NudF, RMSF is found to be within ∼0.5 to 2nm, although its three C-term loop residues and two preceding terminal 11-residue α-helices, connected through a 5-residue loop, shows a stark increment crossing ∼3nm. In contrast, the DMAPP and IPP complexes orderly show the RMSF divergence between ∼0.1-0.8nm and ∼0.06-0.525nm, and here the terminal double-helix does not show any significant displacement. It is interesting to note that the RMSF score across the chain for NudF-IPP is lesser than the NudF-DMAPP scores, indicating that the former is a more stable complex structure.

SASA is another important metric for assessing the extent of receptor exposure to surrounding solvent molecules (Kumar et al. 2021, Ghosh et al. 2020), and as ligand-binding imparts conformational changes in receptor, its respective substructural variations are bound to alter the SASA (Rahman et al. 2020). Although a significant deviation is not observed in the per-residue SASA score, with a comparatively large surface area or a lower compactness of this substructure, as observed earlier, the SASA scores are found to be highly variant across the simulation trajectory. The SASA for NudF structure is observed to consistently drop along the trajectory from 112.5 to 97.5nm^2^, although the respective score of DMAPP complex declines from 117.5 to 97.5nm^2^, with a sharp drop at 20ns. In contrast, the corresponding score for the IPP complex declines from 112.5 to 100nm2, and except for a few time intervals, minute variations were observed throughout the simulation period. SASA for the IPP complex drops dramatically from 110nm2 to 103nm2 after 5ns. The equivalent declination for the DMAPP complex, on the other hand, is noticed after 17.5ns, and its trajectory has significantly more unequal undulations. NudF also exhibits this distinctive declination after 10ns, and here it could indicate a significant functional transition leading to the substantial increase in structural compactness. Moreover, after the initial decline, the SASA graph moves likewise for both DMAPP and IPP complexes.

For estimating the binding affinity of IPP/DMAPP with NudF, the MD trajectories are further investigated to analyze the extent of hydrogen bonds made along the simulation (Lindahl et al. 2010, Abraham et al. 2015). Along the trajectory, NudF shows a nearly 100-120 hydrogen bonds. However, the DMAPP and IPP complexes orderly show 100-130 and 110-130 hydrogen bonds, and a nearly identical variation throughout the trajectory. It indicates their nearly similar scale of atomic interaction within the active site, as also shown by their substantially similar SASA undulations. Although, observing the earlier results, the affinity of DMAPP should not be comparable to that of IPP, consistency of hydrogen bonds is maintained for both the ligand complexes throughout the trajectory, and it indicates the comparable stability of these complexes. To excavate it further, the protein-ligand hydrogen bonding variations are analyzed through the trajectory, and for DMAPP complex, the number of hydrogen bonds is found to slowly increase to two with significantly variant undulations. However, for IPP complex, the number of bonds consistently increases and after ∼60ns, nearly 1 bond is maintained throughout the simulation, showing its more stable interaction. As hydrogen bonds are the major interactions to drive the proper anchoring of ligand within the active site, the higher number of such bonds should be responsible for a stronger interaction (Lindahl et al. 2010, Abraham et al. 2015).

We have observed a few key features. Firstly, the lowest ddG score is found to be the highest for the PHE116 and LYS78 residues, and it confirms that the two residues certainly hold the key to actively evolve the enzyme against the substrates by increasing the structural stability. Secondly, ASP139, GLU140 and ASP141 show a remarkably insignificant ddG score, and it implies that these positions are highly crucial for the NudF function and their top-ranked mutations also failed to stabilize the protein, as earlier discussed by the Nobel Laureate Frances Arnold (Bloom and Arnold, 2009). However, these three residues, along with the substructure T130-L138, show a high RSA score, and it indicates that the flexibility of this superficial loop region could impart a significant functional attribute to NudF. Restricting the search to LYS78 and PHE116 indicates that these positions should be significantly crucial for the stability and interaction affinity of the active site. Thirdly, as LYS78 is superficially more exposed in comparison to ARG116 and is only in contact with one other residues, it should be firstly mutated to study its effect on the overall product yield. It opens venues for their experimental verifications as the top-ranked mutants K78I/K78L and PHE116D/PHE116E could selectively stabilize the conformation and could be responsible for the ligand-specificity, urgently needed to design the novel industrially useful NudF enzyme.

The study extends our understanding about ADP-ribose pyrophosphatase and shows that it has a preferential bias for IPP over DMAPP, with −115.388 (Kcal/mol) versus −41.402 (Kcal/mol) respectively, although if the former is missing, the protein interacts with DMAPP at a much slower rate and probably this could be a key signal to IDI to initiate the conversion of DMAPP to IPP. This promiscuous dephosphorylation of NudF should be studied further through various other substrates, and with that, it would open venues to industrially engineer the cells, wherein the downstream reactions could be channeled more actively with minimal cellular regulation.

## 4. Conclusion

The research thoroughly examines the *Bacillus subtilis* ADP-ribose pyrophosphatase and its 37 functionally confirmed orthologues, and with the projected tertiary structure of the 179-residue representative sequence CUB50584.1, it extensively analyzes the active site of NudF for building knowledgebase for its directed evolution experiments. Although the dataset shows a significantly high phylogenetic divergence, 26.259% residues are found to have a statistically higher evolutionary conservation. According to the findings, ADP-ribose pyrophosphatase has a higher priority interaction with IPP than DMAPP, with −115.388 (Kcal/mol) versus −41.402 (Kcal/mol). The topological variations are restricted to the 8 loop regions, maximally encompassing the active site. Examining the contact map network for the 22 hotspot positions, it is observed that seven residues (ARG18, PRO43, PRO81, ALA117, ASP139, GLU140, and ASP141) do not show even 1 contact and the mutational analysis for the seven prioritized hotspot residues show the highest ΔΔG scores for the LYS78 and PHE116, orderly encoded within loop1 and loop7, encapsulating the active site. The top-ranked mutants F116E, F116D, K78I and K78L show the highest, although equivalent ΔΔG scores of 1.06, 0.97, 0.88, 0.84, and it indicates that these positions should be the key to maximally direct the synthesis of required terpenoids in *Bacillus subtilis*. Thus, the present study must be industrially useful to channelize the entire DXP pathway towards the increased production of prenol or isoprenol or their downstream molecules without generating any metabolic burden.

